# Role of sex and high fat diet in metabolic and hypothalamic disturbances in the 3xTg-AD mouse model of Alzheimer’s disease

**DOI:** 10.1101/2020.07.06.189928

**Authors:** Lisa. S. Robison, Olivia J. Gannon, Melissa A. Thomas, Abigail E. Salinero, Charly Abi-Ghanem, Yannick Poitelon, Sophie Belin, Kristen L. Zuloaga

**Affiliations:** Department of Neuroscience & Experimental Therapeutics, Albany Medical College, Albany, NY, USA 12208

**Author notes:** These authors contributed equally (Lisa. S. Robison, Olivia J. Gannon). Correspondence addressed to: Kristen L. Zuloaga, PhD, Associate Professor, Department of Neuroscience & Experimental Therapeutics, Albany Medical College, 47 New Scotland Avenue; MC-136, Albany, NY, USA 12208.

**Keywords:** sex, Alzheimer’s disease, high fat diet, obesity, inflammation, diabetes, hypothalamus, metabolic

## Abstract

Hypothalamic dysfunction occurs early in the clinical course of Alzheimer’s disease (AD), likely contributing to disturbances in feeding behavior and metabolic function that are often observable years prior to the onset of cognitive symptoms. Late-life weight loss and low BMI are associated with increased risk of dementia and faster progression of disease. However, high fat diet and metabolic disease (e.g. obesity, type 2 diabetes), particularly in mid-life, are associated with increased risk of AD, as well as exacerbated AD pathology and behavioral deficits in animal models. In the current study, we explored possible relationships between hypothalamic function, diet/metabolic status, and AD. Considering the sex bias in AD, with women representing two-thirds of AD patients, we sought to determine whether these relationships vary by sex. WT and 3xTg-AD male and female mice were fed a control (10% fat) or high fat (HF; 60% diet) diet from ~3-7 months of age, then tested for metabolic and hypothalamic disturbances. On control diet, male 3xTg-AD mice displayed decreased body weight, reduced fat mass, hypoleptinemia, and mild systemic inflammation, as well as increased expression of gliosis- and inflammation-related genes in the hypothalamus (Iba1, GFAP, TNF-α, IL-1β). In contrast, female 3xTg-AD mice on control diet displayed metabolic disturbances opposite that of 3xTg-AD males (increased body and fat mass, impaired glucose tolerance). HF diet resulted in expected metabolic alterations across groups (increased body and fat mass; glucose intolerance; increased plasma insulin and leptin, decreased ghrelin; nonalcoholic fatty liver disease-related pathology). HF diet resulted in the greatest weight gain, adiposity, and glucose intolerance in 3xTg-AD females, which were associated with markedly increased hypothalamic expression of GFAP and IL-1β, as well as GFAP labeling in several hypothalamic nuclei that regulate energy balance. In contrast, HF diet increased diabetes markers and systemic inflammation preferentially in AD males but did not exacerbate hypothalamic inflammation in this group. These findings provide further evidence for the roles of hypothalamic and metabolic dysfunction in AD, which in the 3xTg-AD mouse model appears to be dependent on both sex and diet.

## Introduction

Dementia is estimated to affect nearly 40 million people worldwide, with prevalence expected to triple by the year 2050 [1]. Alzheimer’s disease (AD) is the most common form of dementia, with characteristic brain pathology including amyloid beta (Aβ) plaques and tau tangles, that are believed to start forming decades prior to cognitive decline, in addition to inflammation and neurodegeneration. Although cognitive impairment and associated brain regions (e.g. hippocampus, cortex) have been most widely studied, there are numerous non-cognitive symptoms and associated areas of the brain affected that are commonly observed in AD [2]. Further exploration of these phenomena is necessary to prevent and/or treat non-cognitive complications of AD, which may further aggravate disease progression, contribute to poor quality of life, and may even be life-threatening.

One such group of non-cognitive symptoms seen in AD is metabolic disturbances. Late-life weight loss and low body mass index (BMI) are associated with increased dementia risk, faster disease progression, and increased morbidity and mortality [3–5]. In fact, weight loss is a common clinical feature of AD, reported in up to ~40% of cases [6]. Additionally, in both healthy older adults and AD patients, weight loss or low BMI is associated with pathological features of AD, such as increased amyloid burden and CSF biomarkers, as well as cognitive impairment [3, 5, 7–10], suggesting its utility as a biomarker and possible indicator of disease progression. Disturbances in feeding behavior and metabolic function, as well as changes in body weight and composition, are not only seen in AD patients, but also mouse models of the disease. Medications currently approved for AD cannot halt or reverse the progression of the disease, likely because they are begun too late in the disease process. AD patients are usually first diagnosed and begin treatment at the onset of cognitive symptoms, but accumulation of AD pathology in the brain can begin ~20 years earlier. Weight loss is sometimes seen in AD patients during the prodromal phase, which can be more than a decade prior to cognitive decline [11–13], highlighting the possibility of using unintentional weight loss or other metabolic markers as early biomarkers of the disease to identify at-risk individuals.

Metabolic disturbances in AD may be linked to changes in hypothalamic function, which plays a role in regulating energy homeostasis, including feeding behavior. The hypothalamus can be viewed as the control center of the endocrine system, consisting of several distinct nuclei that function to integrate peripheral and central signals. This includes information on body weight/adiposity (insulin and leptin), hunger and satiety signals from the gut (e.g. ghrelin), as well as circulating nutrients and metabolites (e.g. glucose, lipids). The hypothalamus can then signal to other brain regions and the periphery to balance energy intake and expenditure (e.g. control food intake, activity output, thermogenesis) [14]. An accumulation of AD pathology, as well as structural and functional abnormalities, have been noted in the hypothalamus of AD patients, likely contributing to the aforementioned disturbances in energy balance. Both amyloid and tau pathology have been observed in several hypothalamic nuclei of AD patients, including those that regulate energy homeostasis [15]. Neuroimaging studies report decreases in hypothalamic volume/gray matter [16–18], perfusion [19, 20], and glucose metabolism [21] in MCI and/or AD patients. Abnormalities in hypothalamic metabolism and cerebral blood flow have also been seen in a mouse model of AD, observed prior to disturbances in the hippocampus and the onset of cognitive decline [22, 23]. Additionally, previous studies suggest that AD pathology may attenuate central responsivity to peripheral metabolic signals. For example, in the Tg2476 mouse model of amyloid pathology, NPY neurons of the arcuate nucleus in the hypothalamus were found to be less responsive to both leptin and ghrelin treatment, an effect that was mimicked by Aβ treatment in slices from WT mice [24]. It appears that the hypothalamus is not only a target of AD pathology, but that metabolic and hypothalamic dysfunction are also key drivers of disease [2]. Hormones involved in the regulation of energy balance have also been shown to specifically affect amyloid processing, tau, and inflammation [25–30]. Additionally, glucose levels, as well as leptin and insulin signaling, not only play a role in maintaining metabolic homeostasis but are also vital to maintaining hippocampal function and supporting cognitive processes [31–34]. Therefore, interventions targeting metabolic and hypothalamic dysfunction may also contribute to improved outcomes in AD.

In contrast to late-life weight loss, mid-life obesity and metabolic disease (e.g. Type 2 diabetes or prediabetes) are associated with an increased risk for cognitive decline and AD [35, 36]. Metabolic disease and AD are highly comorbid, co-occurring in ~80% of patients [37]. Brains of AD patients commonly exhibit defective insulin signaling [38, 39], leading some to argue that AD is actually “Type 3 diabetes” [40]. Consumption of a hypercaloric diet (e.g. high fat, Western) and metabolic disease have been shown to exacerbate AD pathology and related behaviors in rodent models [41–44]. Mechanistically, metabolic disease also causes dysregulation of hormones involved in energy balance, as well as increased inflammation and amyloidosis, which is common to AD [45–47]. The relationship between metabolic disease and AD may be bidirectional, with AD pathology contributing to metabolic alterations [37, 44, 48]. Despite high rates of comorbidity of metabolic disease in AD patients, little is known about the synergistic effects of AD pathology and high fat diet on the hypothalamus and peripheral markers of metabolic disease. We explored this relationship in the 3xTg-AD mouse model of AD, which exhibits both amyloid and tau pathology, modeling metabolic disease using a chronic high fat diet. We also aimed to determine whether any of these effects were sex-dependent, given that women have greater prevalence of AD and obesity particularly in late life [49, 50].

## Methods

### Animals and experimental design

Male and female 3xTg-AD breeder pairs (#34830-JAX) were obtained from Jackson Laboratories (Bar Harbor, Maine) and were used to breed male and female 3xTg-AD mice for this experiment. These 3xTg-AD mice are on a C57BL/6;129X1/SvJ;129S1/Sv genetic background and exhibit three human mutant genes that result in familial AD, including APPSwe, tauP301L, and Psen1^tm1Mpm^ [51]. Intracellular Aβ deposition can be seen as early as 3-4 months of age, in addition to extracellular Aβ deposition, impaired synaptic transmission and LTP by 6 months, and hippocampal deposits of hyperphosphorylated tau at 12-15 months [51, 52]. Male and female B6129SF2/J mice (#101045) were obtained from Jackson Laboratories (Bar Harbor, Maine) to be used as wild-type (WT) controls. At approximately 3 months of age, mice were put on either a HF diet (60% fat, 5.24 kcal/g; D12492, Research Diets, New Brunswick, NJ) or a control diet (CON; 10% fat; D12450B, Research Diets). This age was chosen, as we previously found that starting high fat diet in young adulthood results in a prediabetic phenotype (weight gain and glucose intolerance) that is similar in male and female WT mice [53]. These mice were part of a larger study and received a sham surgery (~10 min under isoflurane anesthesia, small incision in the neck, sealed with tissue adhesive), followed by buprenex injections 2x daily for 3 days to alleviate pain. The short sham surgery did not cause weight loss in either sex.

Following 3 months on the diet, mice (N=18-25 per group) underwent a glucose tolerance test (GTT) to assess diabetic status. After 4 months on the diet, mice were mice were deeply anesthetized with anesthesia (~4% isoflurane), and blood (cardiac puncture) and tissues were collected. Wet weights were taken of heart, fat pads (visceral and subcutaneous), and reproductive organs (seminal vesicles or uterus). This study was conducted in accordance with the National Institutes of Health guidelines for the care and use of animals in research, and protocols were approved by the Institutional Animal Care and Use Committee at Albany Medical College, Albany, NY, USA.

### Glucose Tolerance Test

At 3 months post-dietary intervention, mice were fasted overnight, and baseline glucose levels were measured by glucometer (Breeze 2, Bayer, Tarrytown, NY). Each mouse received an i.p. injection of 20% glucose at 10 uL/g of body weight, and blood glucose levels were re-measured at 15, 30, 60, 90, and 120 min.

### Open Field Test

An open field test was performed after mice were on respective diets for at least three months. Mice were placed in a square arena and allowed to explore freely for 10 min, then removed and placed in a “recovery cage” so as not to expose them to naïve cage mates. Distance traveled was assessed as a measure of general activity levels.

### Plasma Diabetes Markers & Cytokines

Mice had been fasted ~5 hours prior to euthanasia and blood collection. During euthanasia, blood was collected via cardiac puncture, mixed with 5uL of EDTA on ice, and spun at 1,500 x g for 10 minutes at 4°C to collect plasma. Plasma samples were stored at −80°C until assayed. Diabetes-associated markers and cytokine levels in the plasma were assessed using the Bio-Plex Pro Mouse Diabetes 8-Plex Assay (Cat# 171F7001M; Bio-Rad, Carlsbad, California) and the Bio-Plex Pro Mouse Cytokine 23-Plex Assay (Cat# M60009RDPD; Bio-Rad, Carlsbad, California), respectively, according to the manufacturer’s instructions.

### Liver Histology

After livers were removed from mice during euthanasia, they were post-fixed for 24 hours in 4% formalin, then placed in 30% sucrose solution for 72 hours. Livers were frozen in OCT compound and stored at −80°C until being cut at 5 μm and stained with either Hematoxylin and Eosin (H&E) or Sirius Red.

#### H&E Staining and Analysis

Hallmarks of NAFLD, steatosis, ballooning, and inflammation, were assessed using hematoxylin and eosin (H&E) staining. Sections were equilibrated to room temperature before being washed in Phosphate Saline Buffer (PBS) for five minutes. The sections were stained in 0.1% Mayer’s Hematoxylin (Sigma MHS-16) for 10 minutes and 0.5% Eosin (Sigma Eosin Y-solution 0.5% alcoholic) for 10 minutes. They were dipped in 50% EtOH ten times, 70% EtOH ten times, equilibrated in 95% EtOH for 30 seconds and in 100% EtOH for 1 minute. The sections were then dipped in xylene multiple times, mounted with Cytoseal XYL (ThermoFisher 8312-4), and cover slipped. Sections were viewed on a Zeiss Primo Vert Inverted Phase Contrast microscope and imaged with an attached Axiocam 105 color camera.

The NAFLD activity score is a semi-quantitative method of analyzing the hallmarks of NAFLD, including steatosis, ballooning, and inflammation. These three attributes characterize the disease and each category is graded on a scale of 0-3, adapted from a previous study [54]. Steatosis, or microvesicular fat, is defined as faint or patchy white areas within cell cytoplasm. A score of 0 indicates no steatosis while a score of 3 indicates mostly steatosis. Ballooning is defined as larger, circular aggregates of fat between cells (macrovesicular fat). A score of 0 indicates no ballooning while a score of 3 indicates the greatest amount of ballooning. Inflammation is characterized by the clustering of leukocytes resulting in focal inflammation. A score of 0 is no inflammation while a score of 3 is distinctive clusters. Three liver sections per animal were rated to obtain this 3-part score (0-3 for steatosis, ballooning, and inflammation).

#### Sirius Red Staining and Analysis

Sections were equilibrated to room temperature before being incubated in 0.1% Sirius Red (Sigma “Direct Red 80”) for 1 hour. The sections were then washed in acidified water twice and dipped in 100% EtOH three times. They were cleared in xylene and mounted with Cytoseal XYL. The sections were viewed on a Zeiss Primo Vert Inverted Phase Contrast microscope and imaged with an attached Axiocam 105 color camera. In ImageJ (NIH), images were converted into 8-bit files and underwent thresholding. Regions of interest were drawn on liver sections, and fibrosis was measured as percent area positive for staining.

### Immunofluorescence

Mice used for immunofluorescence were transcardially perfused with 0.9% saline. Brains were rapidly removed and post-fixed overnight in 4% formalin, then cryoprotected in 30% sucrose for at least 72h, embedded in OCT, and stored at −80°C until cryosectioning. Brains were sectioned in the coronal plane into 6 series of 40 uM thick sections. Sections containing the hypothalamus were washed with PBS with 0.01% sodium azide, permeabilized in 0.3% TPBS with sodium azide for 1h at room temperature, blocked in 4% donkey serum in 0.3% TPBS with sodium azide for 1h at room temperature, and incubated at 4°C overnight with primary antibodies. Primary antibodies for the evaluation of microglia/macrophages included goat anti-Iba1 (1:1000, PA5-18039, ThermoFisher, Lot # TI2638761), while rat anti-glial fibrillary acidic protein (1:2500, AB5804, Millipore, Lot # TA265137) was used for the evaluation of astrocytes. Rabbit anti-NeuN (1:1000, ABN78, Millipore, Lot # 3041797) or DAPI (1:1,000; added with secondary antibodies) were used to visualize hypothalamic nuclei and draw regions of interest for analysis (see next section). Fluorescent secondary antibodies (Jackson ImmunoResearch, West Grove, PA), including Rhodamine Red-X Donkey Anti-Rabbit (1:100), Alexa Fluor 647 Donkey Anti-Goat (1:300), and DyLight™ 405 AffiniPure Donkey Anti-Rat (1:300), were diluted in blocking buffer and applied at room temperature for 2h.

### Immunofluorescence analysis

Images for quantification were taken of the hypothalamus at 10x using the Axio Observer fluorescent microscope (Carl Zeiss Microscopy, Jena, Germany). All immunohistochemistry analyses were performed in coronal sections across the hypothalamus. All measurements were performed by an experimenter who was blinded to the identity of the treatment group from which sections came. Iba1 and GFAP images were separately thresholded using ImageJ (NIH, Bethesda, MD, USA) software. NeuN labeling or DAPI staining were used to visualize hypothalamic nuclei and draw regions of interest. Regions of interest were drawn around the arcuate nucleus (ARC), dorsomedial hypothalamus (DMH), ventromedial hypothalamus (VMH), lateral hypothalamus area (LHA) and paraventricular nucleus (PVN) of every 6^th^ section using ImageJ to quantify the average area covered by cells positive for each of these antibodies.

### Quantitative reverse transcriptase-PCR (RT-qPCR)

During euthanasia, brains were rapidly removed and hypothalamus microdissected in PBS over dry ice, then stored at −80°C until RNA isolation was performed. Total RNA was extracted using the TRIZOL method. All samples containing RNA were treated using the TURBO DNA-free Kit (Invitrogen, Catalog number: AM1907) before reverse transcription. cDNA was prepared using 1 μg RNA and the High-Capacity cDNA Reverse Transcription Kit (Applied Biosystems, Catalog number: 4368814). The qPCR reactions were performed using TaqMan Gene Expression Master Mix (Applied Biosystems, Catalog number: 4369016) in the presence of Taqman Assays with primer/probes for Iba1 (Mm00479862_g1), GFAP (Mm01253033_m1), IL-1β (Mm00434228_m1), TNF-α (Mm00443258_m1), IL-6 (Mm00446190_m1), AgRP (Mm00475829_g1), NPY (Mm01410146_m1), POMC (Mm00435874_m1), LepR (Mm00440181_m1), MCR4 (Mm00457483_s1), and FNDC5 (Mm01181543_m1) as target genes, and HPRT (Mm01324427_m1), RPL13A (Mm01612986_gH), and RPS17 (Mm01314921_g1) as housekeeping genes. Samples, housekeeping genes, water (negative control), and a reference/positive control sample were run in triplicate. Statistical analyses were performed on ΔCt values, and data was plotted as relative normalized expression compared to male WT control fed mice using CFX Maestro Software (Bio-Rad Laboratories, Inc., Hercules, CA, USA).

### Statistical Analysis

All data are expressed as mean + SEM. Four-way repeated measures ANOVAs (between-subjects measures: AD, sex, diet; within-subject measure: time) were used for analyses of monthly body weight and blood glucose levels over time. Analyses of all other measures were analyzed using a three-way ANOVA (between-subjects measures: AD, sex, diet). ANOVA’s were followed by posthoc tests (Tukey method) when appropriate. Correlations were run separately for each sex/genotype (WT males, AD males, WT females, AD females) to assess the relationships between metabolic and hypothalamic abnormalities. Statistical significance was set at p<0.05, and statistical analyses were performed using GraphPad Prism v8, SigmaStat v12, or Statistica v13 software.

## Results

### Sex and diet interact to influence weight gain, adiposity, and glucose intolerance in 3xTg-AD mice

#### Weight gain and adiposity

To assess sex differences in the metabolic effects of high fat diet, we measured body weight monthly **(Figure 1A)** and calculated weight gain throughout the entire diet intervention **(Figure 1B).** Adiposity was assessed by measuring the mass of subcutaneous **(Figure 1C)** and visceral **(Figure 1D)** fat wet weights upon euthanasia. On control diet, AD males weighed less than WT males (month 2 through the end of the experiment; p<0.05 for all), while the opposite was seen in females, with AD female mice weighing more than WT female mice (month 2, month 3, end weight; p<0.05 for all). On control diet, AD males also had less subcutaneous fat and visceral fat than both WT males (p<0.001) and AD females (p≤0.001). In all groups, mice on a HF diet gained significantly higher percentage of weight (p<0.0001 for all), subcutaneous fat (p<0.0001 for all), and visceral fat (p<0.001 for all) compared to their control diet fed counterparts. Of note, AD HF fed females gained a higher percentage of weight (p<0.0001 vs. WT HF females and AD HF males), and had more subcutaneous fat (p<0.0001 vs. WT HF females and AD HF males) and visceral fat (p<0.0001 vs. all).

**Figure 1:**
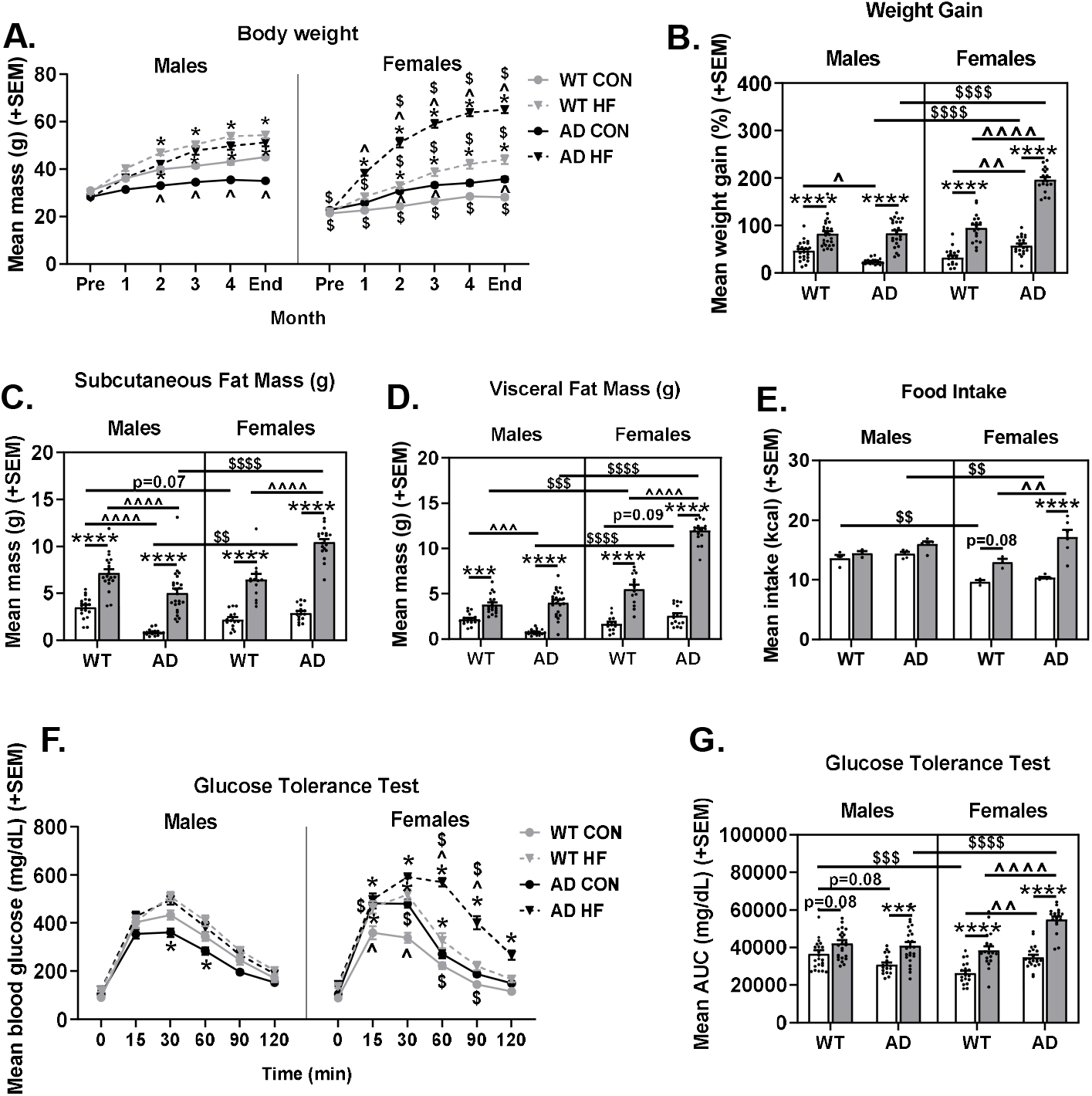
Sex and diet interact to influence weight gain, adiposity, and glucose intolerance in 3xTg-AD mice. Male and female B6129SF2/J mice (WT) and 3xTg-AD mice were placed on control (CON; 10% fat) or high fat (HF; 60% fat) diet at ~3 months of age, which was continued for the duration of the experiment. After ~3 months on the diet, mice were subjected to glucose tolerance testing (GTT). Tissue collection was performed after ~4 months on respective diet, at ~7 months of age. (A) Body weight was measured at the beginning of the experiment, prior to the start of respective diet intervention, once per month during the diet intervention, and at the end of the experiment just prior to tissue collection. N=18-25/group. (B) Weight gain was calculated as the percent difference in body weight at the end of the experiment versus initial body weight measured prior to the start of diet intervention. N=18-25/group. (C) Subcutaneous and (D) visceral fat wet weights (in grams) assessed at the end of the experiment. N=14-24/group. (E) Food intake was measured (in grams) to obtain an average measure of daily food intake for each cage of mice within the same treatment group. The mass of food consumed was multiplied by the energy density in each respective diet type to obtain the average daily caloric intake. Each data point represents the mean intake of one cage of mice (3-5 mice per cage). N=3-6 cages/group. (F) Glucose tolerance testing (GTT). GTT was performed to assess diabetic status 3 months after the start of diet intervention. Blood glucose levels were measure following overnight fasting (t=0), then mice were injected with glucose challenge and blood glucose levels were measured at 15, 30, 60, 90, and 120 min post-injection. (G) Area under the curve (AUC) was calculated. N=18-25/group. * = diet effect p<0.05; ** = diet effect p<0.01; *** = diet effect p<0.001; **** = diet effect p<0.0001 $ = sex effect p<0.05; $$ = sex effect p<0.01; $$$ = sex effect p<0.001; $$$$ = sex effect p<0.0001 ^ = AD effect p<0.05; ^^ = AD effect p<0.01; ^^^ = AD effect p<0.001; ^^^^ = AD effect p<0.0001

Some of the observed differences in body weight and adiposity could be related to changes in energy intake and expenditure; therefore, food intake **(Figure 1E)** and activity levels in an open field test were assessed. WT (p=0.0781) and AD (p<0.0001) females, but not males, on a HF diet consumed more food compared to their control diet fed counterparts. Additionally, AD HF females ate more than WT HF females (p=0.0035). The open field test revealed that reduced body mass and adiposity observed in AD males compared to WT males was not associated with increased locomotor behavior (p>0.05); in fact, AD HF males were hypoactive compared to WT HF males (p=0.0208). However, increased weight gain and adiposity in AD HF females was indeed associated with reduced activity levels in the open field compared to AD LF females (p=0.0018). Means ± SEM of distance traveled in the open field (m) were WT M CON: 12.368 ± 2.336, WT M HF: 19.383 ± 2.414, AD M CON: 8.843 ± 2.220, AD M HF: 9.339 ± 1.867, WT F CON: 22.539 ± 2.477, WT F HF: 20.068 ± 2.140, AD F CON: 28.174 ± 2.874, AD F HF: 15.023 ± 2.111.

#### Organ mass

Heart mass was assessed as a surrogate measure of cardiac stress, as it is increased by obesity [54, 55]. The HF diet also increased heart mass in AD females (p=0.0235 vs. LF AD Female, p=0.0002 vs. HF WT female), but not males. Means ± SEM of heart mass (g) were WT M CON: 0.221 ± 0.006, WT M HF: 0.220 ± 0.009, AD M CON: 0.229 ± 0.004, AD M HF: 0.238 ± 0.008, WT F CON: 0.147 ± 0.006, WT F HF: 0.143 ± 0.005, AD F CON: 0.156 ± 0.005, AD F HF: 0.189 ± 0.007.

Reproductive organ weight was assessed as a surrogate measure of sex hormone levels [55–57]. Neither AD nor diet significantly affected reproductive organ mass (p>0.10 for all) in either sex, suggesting that it is unlikely that hormone levels were altered by these treatments. Means ± SEM of reproductive organ mass (g) were WT M CON: 0.430 ± 0.022, WT M HF: 0.451 ± 0.019, AD M CON: 0.424 ± 0.019, AD M HF: 0.448 ± 0.019, WT F CON: 0.180 ± 0.015, WT F HF: 0.187 ± 0.015, AD F CON: 0.240 ± 0.020, AD F HF: 0.206 ± 0.014.

#### Glucose tolerance test

A glucose tolerance test was performed after approximately 3 months on respective diets. Blood glucose levels were assessed immediately prior to and at intervals of up to two hours after a glucose challenge injection **(Figure 1F).** Analysis of area under the curve (AUC) following the glucose challenge was also computed **(Figure 1G)**. On a control diet, WT females had better glucose tolerance compared to males (p=0.0003). Additionally, on control diet, AD males trended towards having better glucose tolerance compared to WT counterparts (p=0.081). Conversely, AD females had impaired glucose tolerance compared to WT counterparts (p=0.0053). HF diet impaired glucose tolerance in all groups; however, this did not reach significance for WT males (p=0.081; p<0.001 for all others). Of note, HF diet impaired glucose tolerance to the greatest degree in AD females (p<0.0001 vs. WT HF females and AD HF males).

### HF diet and AD are associated with NAFLD pathology

Nonalcoholic fatty liver disease (NAFLD) is a condition in which excess fat accumulates in the liver. It is the most common cause of liver disease in the U.S., affecting ~30% of adults [58]. Nonalcoholic steatohepatitis (NASH) is a form of NAFLD that also includes inflammation (infiltration of inflammatory immune cells and secretion of pro-inflammatory cytokines) and liver cell damage that can progress to fibrosis (scarring), cirrhosis, and/or liver cancer [59]. We evaluated the presence and severity of NAFLD in these mice not only because of the interrelationship between metabolic disease and NAFLD [60], but also because evidence suggests that NAFLD may contribute to AD [61].

NAFLD was assessed by scoring liver sections stained with hematoxylin and eosin (H&E; **Figure 2E**) for steatosis (intracellular microvesicular fat; **Figure 2A**), ballooning (extracellular macrovesicular fat; **Figure 2B**), and inflammation (leukocyte accumulation; **Figure 2C**). Compared to males, females had greater steatosis (trend, p=0.0668) and ballooning (p<0.0001); this sex difference was particularly pronounced in mice on control diet. Conversely, females had lower inflammation scores overall (p<0.0001). As expected, HF diet increased steatosis (p=0.0001) and inflammation (p<0.0001), though HF diet increased ballooning in males only (sex x diet interaction p=0.0011). While there were no differences between WT and AD mice in steatosis, AD mice had greater ballooning (p<0.0001) and inflammation (p<0.0001) compared to WT mice. AD males also appeared to be most susceptible to HF diet-induced NAFLD pathology, including ballooning (p=0.0193 vs. WT HF males) and particularly inflammation (p<0.05 vs. all other groups).

**Figure 2.**
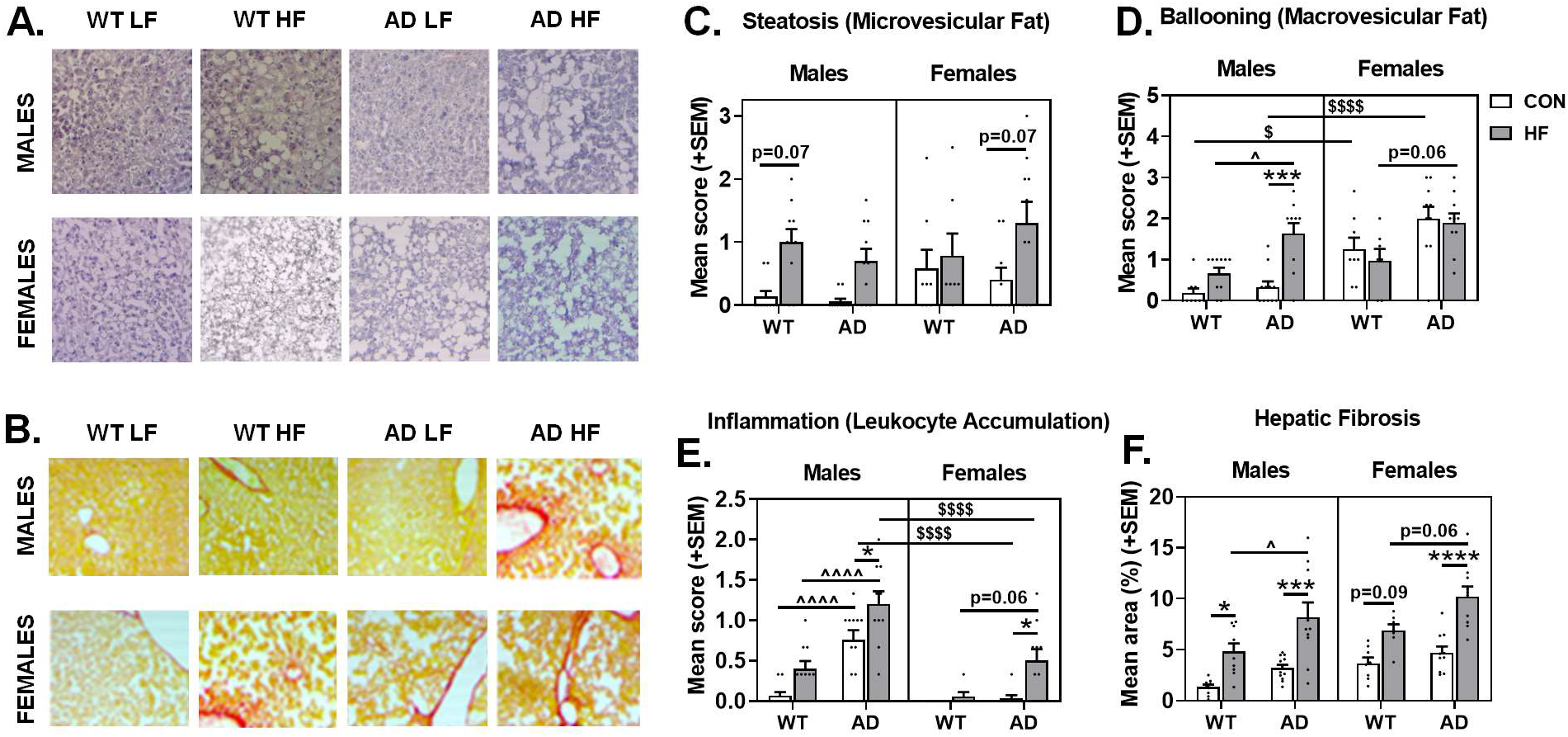
HF diet and AD are associated with nonalcoholic fatty liver disease (NAFLD)-related pathology. Liver sections were stained with (A) hematoxylin and eosin (H&E) or (B) Sirius red to assess NAFLD pathology. H&E stained sections were assessed for (C) steatosis (microvesicular fat), (D) ballooning (macrovesicular fat), and (E) inflammation (leukocyte accumulation) using a semi-quantitive scoring system (scored 0-3). (F) Sirius red stained sections were assessed for hepatic fibrosis by measuring the % area positive for the stain. N=7-12/group for all measures. * = diet effect p<0.05; ** = diet effect p<0.01; *** = diet effect p<0.001; **** = diet effect p<0.0001 $ = sex effect p<0.05; $$ = sex effect p<0.01; $$$ = sex effect p<0.001; $$$$ = sex effect p<0.0001 ^ = AD effect p<0.05; ^^ = AD effect p<0.01; ^^^ = AD effect p<0.001; ^^^^ = AD effect p<0.0001

Hepatic fibrosis was assessed by %area positive for Sirius Red stain **(Figure 2D, 2F)**. As expected, HF diet increased fibrosis (main effect of diet; p<0.0001). Additionally, AD mice had greater fibrosis compared to WT mice (main effect of AD, p<0.0001), and females had greater fibrosis than males (p=0.0009).

### Sex-specific effects of HF diet and AD on plasma levels of diabetes-associated markers

Plasma was assayed for diabetes-associated markers **(Figure 3A-H)**. Leptin **(Figure 3B)** and ghrelin **(Figure 3C)** are hormones that have opposing actions on energy balance via signaling in the hypothalamus. Leptin is primarily secreted by adipocytes to attenuate food intake and promote energy expenditure, serving as a mediator of long-term energy balance regulation [62]. Of note, we found that AD males on control diet exhibited severe hypoleptinemia, such that plasma leptin levels were barely detectable. As expected, HF diet resulted in hyperleptinemia (p<0.0001 main effect of diet). This diet effect was significant in WT males (p=0.0141), AD males (p<0.0001), and AD females (p=0.0006), but not WT females. In females on HF diet, AD mice had higher leptin levels than WT mice (p=0.0326). Ghrelin is a fast-acting anorexigenic hormone produced in the gut, serving to stimulate feeding behavior [62]. Mice on HF diet had lower ghrelin levels compared to control diet fed mice (p<0.0001 main effect of diet). Within WT mice, females had higher levels of ghrelin than males (p=0.019), while AD males had higher ghrelin levels than WT males (p=0.015).

**Figure 3.**
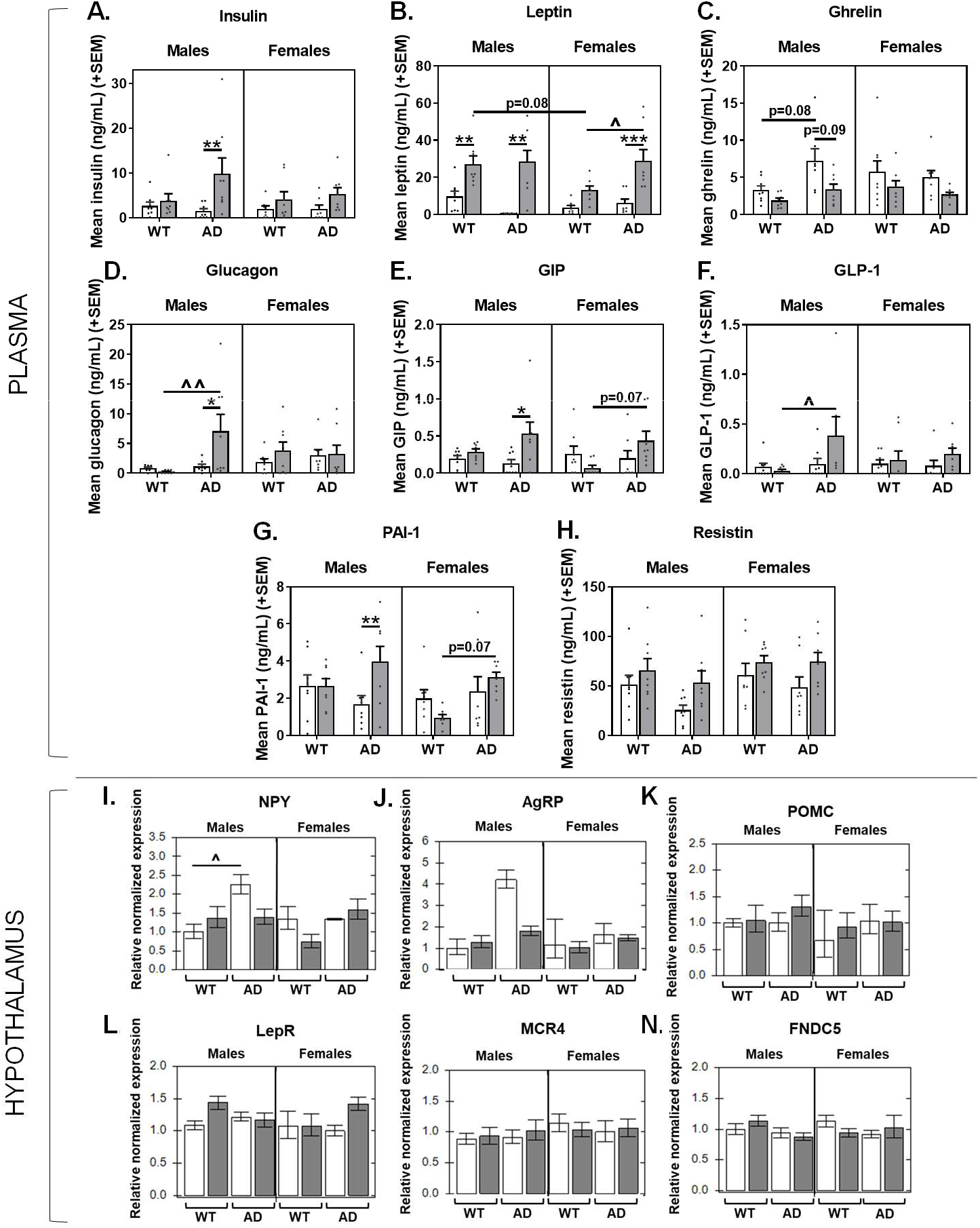
Sex-specific effects of HF diet and AD on plasma levels of diabetes-associated markers and expression of hypothalamic peptides that regulate feeding. (A-H) Plasma concentrations of diabetes-related markers. Blood was collected following a 5 hour fasting period just prior to euthanasia, and assayed for (A) insulin, (B) leptin, (C) ghrelin, (D) glucagon, (E) gastric inhibitory polypeptide (GIP), (F) glucagon-like-peptide 1 (GLP-1), (G) plasma plasminogen activator inhibitor-1 (PAI-1), and (H) resistin. N=8/group for all plasma markers. (I-N) Gene expression levels in homogenate of the whole hypothalamus related to energy balance. Hypothalamus was collected following a 5 hour fasting period just prior to euthanasia, and assayed for (I) neuropeptide Y (NPY), (J) agouti-related peptide (AgRP), (K) pro-opiomelanocortin (POMC), (L) leptin receptor (LepR), (M) melanocortin receptor 4 (MCR4), (N) Fibronectin type III domain-containing protein 5 (FNDC5). N=5-8/group for all gene expression analyses in the hypothalamus. * = diet effect p<0.05; ** = diet effect p<0.01; *** = diet effect p<0.001; **** = diet effect p<0.0001; $ = sex effect p<0.05; $$ = sex effect p<0.01; $$$ = sex effect p<0.001; $$$$ = sex effect p<0.0001; ^ = AD effect p<0.05; ^^ = AD effect p<0.01; ^^^ = AD effect p<0.001; ^^^^ = AD effect p<0.0001

Insulin **(Figure 3A)** and glucagon **(Figure 3D)** are hormones involved in maintaining homeostatic blood glucose levels, secreted by pancreatic β-cells and α-cells of the islet of Langerhans, respectively. While insulin promotes cells absorbing glucose, thus reducing blood glucose levels to prevent hyperglycemia, glucagon signals for the release of stored glucose from the liver, resulting in increased blood glucose levels to prevent hypoglycemia [63]. HF diet resulted in hyperinsulinemia in AD males (p=0.0092). AD HF males also had higher levels of glucagon compared to AD control fed males (p=0.0399) and WT HF males (p=0.0080).

Gastric inhibitory polypeptide (GIP; **Figure 3E**) and glucagon-like-peptide 1 (GLP-1; **Figure 3F**) and are two incretin hormones secreted by cells in the intestine that promote insulin secretion. However, they have opposing actions on postprandial glucagon secretion, such that GIP enhances it and GLP-1 suppresses it. GLP-1 and GIP also act in the hypothalamus to induce satiety and decrease food consumption [64]. AD HF males had higher GLP-1 levels compared to WT HF males (p=0.0363). Levels of circulating GIP also appeared to be highest in AD HF males. Though mice on HF diet had higher levels of GIP compared to control diet fed mice (p<0.0001 main effect of diet), this was driven by AD HF males which had higher GIP levels compared to their control fed counterparts (p=0.0421), and AD HF females tended to have higher levels than WT HF females (p=0.0743).

Plasma plasminogen activator inhibitor-1 (PAI-1; **Figure 3G**) is synthesized by adipose tissue and is responsible for the inhibition of tissue plasminogen activator and urokinase, enzymes that convert plasminogen to plasmin. Plasmin plays a key role in fibrinolysis, degrading fibrin and extracellular matrix proteins to protect against pathological fibrin deposition and tissue damage. Increases in PAI-1 can result from inflammatory signals, and it can contribute to a pro-thrombotic state and the development of metabolic disease [65]. HF AD mice had higher levels than WT mice (p=0.002). AD HF males had higher levels than AD control fed males, while AD HF females had higher levels than WT HF females (p=0.0705).

Resistin **(Figure 3H)** is a hormone secreted by adipose tissue, as well as immune and epithelial cells, which suppresses insulin’s ability to promote cellular glucose uptake. Resistin has been suggested to be an important link between obesity, insulin resistance and diabetes. Mice on HF diet had higher levels of resistin compared to control diet fed mice (p=0.0050 main effect of diet). Although not significant, this tended to be more pronounced in AD compared to WT mice.

### Hypothalamic expression of orexigenic peptides is increased in AD males on control diet

Several signaling molecules work together in the hypothalamus to regulate energy homeostasis, balancing food consumption with energy expenditure. Relative expression of genes involved in energy balance were assessed in homogenate of the whole hypothalamus **(Figure 3I-N)**. This included neuropeptide Y (NPY; fast-acting orexigenic peptide; **Figure 3I**) and agouti-related peptide (AgRP; delayed, longer-acting orexigenic peptide; **Figure 3J**) [66], Pro-opiomelanocortin [POMC; an anorexigenic precursor protein and member of the central melanocortin system; cleaved to α-melanocyte stimulating hormone (α-MSH)] [67] **(Figure 3K)**, leptin receptor (LepR; binds the anorexigenic hormone, leptin; **Figure 3L**), melanocortin receptor 4 (MCR4; a G protein-coupled receptor that binds α-MSH; **Figure 3M**), and Fibronectin type III domain-containing protein 5 (FNDC5; the precursor of irisin, a thermogenic adipomyokine**; Figure 3N**) [68].

There was a main effect of AD to increase expression of both NPY (p=0.0058) and AgRP (p=0.0265) compared to WT mice. This effect seems to be primarily driven by control diet fed AD males expressing higher levels of each gene compared to their WT counterparts, though this reached statistical significance for NPY (p=0.0394; AD x sex x diet interaction p=0.0072) but not for AgRP (p=0.1093). There were no other main effects, interactions, or group differences for hypothalamic gene expression levels of POMC, MCR4, LepR, or FNDC5 (p>0.05 for all).

### Systemic inflammation is increased in AD males only and exacerbated by HF diet

Plasma levels of 23 cytokines were assessed as markers for systemic inflammation. Significant group differences were seen in levels of IL-10, IL-12 (p40), MIP-1α, and MIP-1β **(Figure 4A-D)**. There were increases in IL-10 and IL-12 (p40) in AD mice (p<0.01 for both); this main effect was driven by males and exacerbated by HF diet. Similar trends of increased cytokine levels in HF fed AD males were seen for MIP-1α and MIP-1β (p<0.05 vs. all other groups).

**Figure 4.**
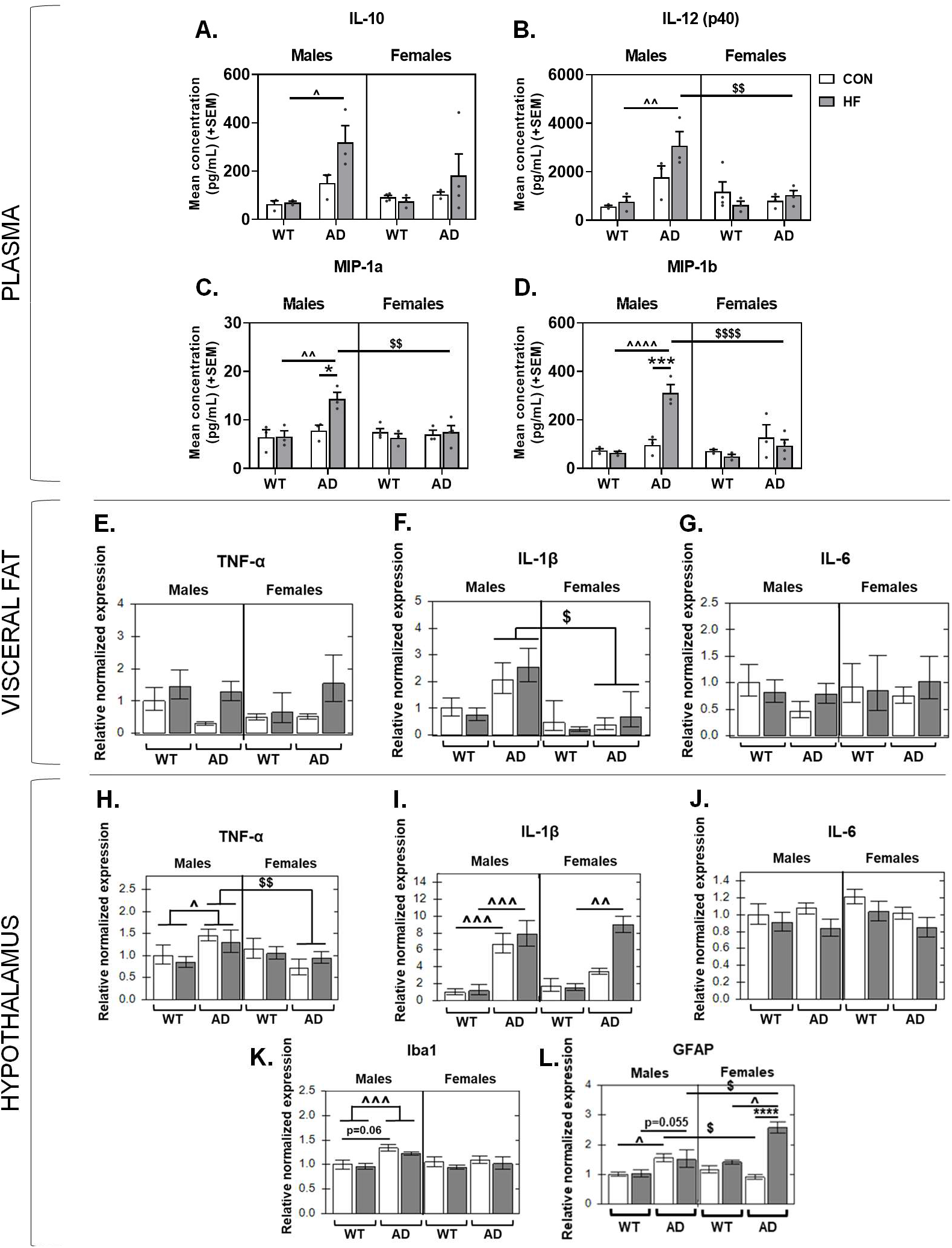
Sex differences in the effects of AD and diet on peripheral inflammation and hypothalamic expression of inflammation-related genes. (A-H) Plasma concentrations of cytokines. Blood was collected following a 5 hour fasting period just prior to euthanasia, and assayed for 23 cytokines, with significant group differences seen for (A) IL-10, (B) IL-12 (p40), (C) MIP-1a, (D) MIP-1b. N=3-4/group for all plasma markers. (E-G) Gene expression levels in visceral fat related to inflammation, including (E) TNF-α, (F) IL-1β, (G) IL-6. (H-L) Gene expression levels in homogenate of the whole hypothalamus related to inflammation. Hypothalamus was collected following a 5-hour fasting period just prior to euthanasia, and assayed for (H) TNF-α, (I) IL-1β, (J) IL-6, (K) Iba1, (L) GFAP. N=5-8/group for all gene expression analyses in the hypothalamus. * = diet effect p<0.05; ** = diet effect p<0.01; *** = diet effect p<0.001; **** = diet effect p<0.0001; $ = sex effect p<0.05; $$ = sex effect p<0.01; $$$ = sex effect p<0.001; $$$$ = sex effect p<0.0001; ^ = AD effect p<0.05; ^^ = AD effect p<0.01; ^^^ = AD effect p<0.001; ^^^^ = AD effect p<0.0001

Visceral fat was also investigated as a potential source of inflammation in the periphery **(Figure 4E-G)**. Notably, gene expression levels of TNF-α were significantly increased by HF diet in AD mice (p=0.0018) but not WT mice. Additionally, AD males had greater IL-1β expression in visceral fat compared to AD females (p=0.0372). Gene expression levels of IL-6 were not significantly different between groups.

### Sex differences in the effects of AD and diet on hypothalamic expression of inflammation-related genes

Neuroinflammation is increased in response to HF diet and is believed to contribute to both metabolic disease and neurodegenerative processes in AD [69, 70]. Relative gene expression for markers of neuroinflammation were assessed in homogenate of hypothalamic tissue, including Iba1, GFAP, IL-1β, TNF-a, and IL-6 **(Figure 4H-L)**.

Ionized calcium-binding adaptor molecule 1 (Iba1) is a calcium-binding protein expressed in monocyte lineage cells such as macrophages, including microglia in the brain. While Iba1 is expressed in both resting and activated microglia [71], it is believed to increase with microglia activation due to its role in membrane ruffling and phagocytosis [72, 73]. Therefore, an increase in expression could be due to an increase in the number of microglia and/or their activation status. AD mice had higher expression of Iba1 compared to WT mice (main effect of AD p=0.0058). This effect seemed to be driven by males (AD x sex interaction p=0.0351), and AD males had higher expression of Iba1 compared to WT males (p<0.001) and AD females (p=0.014). Additionally, AD males on control diet had a trend toward greater Iba1 expression compared to their WT counterparts (p=0.0603).

Glial fibrillary acidic protein (GFAP) is an intermediate filament protein, and its expression is increased in reactive astrocytes and indicative of inflammation [74]. AD males on both control (p=0.0272) and HF (p=0.0558) diet had higher GFAP expression compared to their WT counterparts. AD control males also had greater GFAP expression compared to AD control fed females (p=0.0272). In females, the combination of AD and HF diet resulted in significantly increased GFAP expression compared to all other groups (p<0.05 for all).

TNF-α, IL-1β, and IL-6 are cytokines that have been implicated in both metabolic disease and AD pathogenesis [75–77]. These cytokines are produced by cell in the brain and are also secreted in the periphery, such as in adipose tissue, particularly under obesigenic conditions [78–80]. Mimicking Iba1 results, male AD mice had higher expression compared to WT males (p=0.020) and AD females (p=0.006). Similarly, AD males, regardless of diet, had higher IL-1β expression compared to WT males (p<0.001 for both). AD females only had increased IL-1β expression compared to WT females when on a HF diet (p=0.0020). Lastly, mice on a HF diet had lower expression of IL-6 (p=0.0290 main effect of diet) compared to control fed mice.

Correlations were run to test the hypothesis that changes in inflammation-related gene expression in the hypothalamus were associated with metabolic disturbances **(Table 1)**. In AD males only, weight gain was inversely associated with IL-6 (p<0.05 for all). In AD females only, weight gain was positively associated with IL-1B and GFAP expression (p<0.05 for all). To explore possible mechanisms of hypothalamic astrogliosis in AD females that are associated with weight gain, we ran correlations between GFAP expression with known astrocyte activators, leptin (plasma) and IL-1β (hypothalamic expression) **(Table 1)**. These correlations were run for all groups, but only in AD females was GFAP expression positively associated with both leptin and IL-1β.

**Table 1.** Sex- and genotype- specific correlations between metabolic outcomes and hypothalamic abnormalities. Correlations were run for each sex/genotype (WT males, AD males, WT females, AD females), combining mice across control and high fat diets, to explore relationships between metabolic outcomes and hypothalamic abnormalities.

|  | WT MALES |  | AD MALES |  | WT FEMALES |  | AD FEMALES |  |
| --- | --- | --- | --- | --- | --- | --- | --- | --- |
|  | R | p | R | p | R | p | R | p |
| Weight gain vs. IL-6 (hyp) | -0.049 | 0.863 | **-0.587** | **0.017\*** | -0.201 | 0.473 | -0.395 | 0.230 |
| Weight gain vs. IL-1β (hyp) | 0.201 | 0.491 | 0.280 | 0.312 | -0.401 | 0.155 | **0.806** | **0.0012\*\*** |
| Weight gain vs. GFAP (hyp) | -0.231 | 0.408 | -0.164 | 0.544 | 0.217 | 0.456 | **0.891** | **<0.0001\*\*\*\*** |
| Leptin (plasma) vs. GFAP (hyp) | -0.239 | 0.454 | -0.333 | 0.267 | 0.182 | 0.593 | **0.756** | **0.030\*** |
| IL-1β (hyp) vs. GFAP (hyp) | 0.004 | 0.988 | 0.043 | 0.878 | 0.432 | 0.140 | **0.935** | **<0.0001\*\*\*\*** |
| Weight gain vs. Iba1 (PVN) | 0.031 | 0.933 | 0.704 | 0.051 | -0.069 | 0.870 | **0.775** | **0.009\*\*** |
| Weight gain vs. GFAP (ARC) | -0.022 | 0.956 | -0.005 | 0.992 | 0.490 | 0.151 | **0.780** | **0.008\*\*** |
| Weight gain vs. GFAP (DMH) | **0.759** | **0.018\*** | **0.755** | **0.030\*** | **0.854** | **0.002\*\*** | **0.932** | **<0.0001\*\*\*\*** |
| Weight gain vs. GFAP (VMH) | 0.237 | 0.510 | 0.389 | 0.341 | 0.501 | 0.140 | **0.834** | **0.003\*\*** |
| Weight gain vs. GFAP (PVN) | -0.169 | 0.663 | 0.357 | 0.432 | -0.074 | 0.839 | **0.857** | **0.003\*\*** |

### Sex differences in hypothalamic gliosis in response to AD and HF diet

Since gene expression for Iba1 and GFAP were increased in the homogenate of the whole hypothalamus, immunofluorescence was performed to localize specific nuclei that experience gliosis in response to AD and HF diet. Several nuclei in the hypothalamus work together to regulate energy balance, balancing food consumption with energy expenditure. First order neurons that receive peripheral metabolic signals such as insulin, leptin and ghrelin, in addition to nutrients and metabolites like glucose and fatty acids, lie within the arcuate nucleus (ARC; infundibular nucleus in humans). The two major cells types in the arcuate nucleus express either NPY/AgRP (orexigenic) or POMC/cocaine- and amphetamine-regulated transcript (CART) (anorexigenic). AgRP/NPY and POMC/CART project to other hypothalamic nuclei to regulate food intake and energy expenditure (e.g. locomotor behavior, thermogenesis). This includes the dorsomedial hypothalamus (DMH), lateral hypothalamus area (LHA; major site for hunger) and the ventromedial hypothalamus (VMH; satiety center). Additionally, neurons from the arcuate nucleus project to the paraventricular nucleus (PVN; a major anorexigenic site), which subsequently regulates food intake, as well as energy expenditure via projections to the nucleus tractus solitarius (NTS) in the brainstem [81]. In addition to contributing to inflammation, glial cells have also demonstrated a variety of ways in which they control energy balance [82].

#### Iba1 labeling

Immunolabeling was performed for Iba1 to assess microgliosis in nuclei of the hypothalamus that regulate energy balance **(Figure 5A-F)**. Iba1 immunoreactivity was increased in AD mice in the ARC (p<0.0001), DMH (p<0.0001), and VMH (p=0.0040). Additionally, HF diet increased Iba1 labeling in the PVN of AD mice, but not WT mice (AD x diet interaction p=0.0271). No group differences were seen in Iba1 labeling in the LHA.

**Figure 5.**
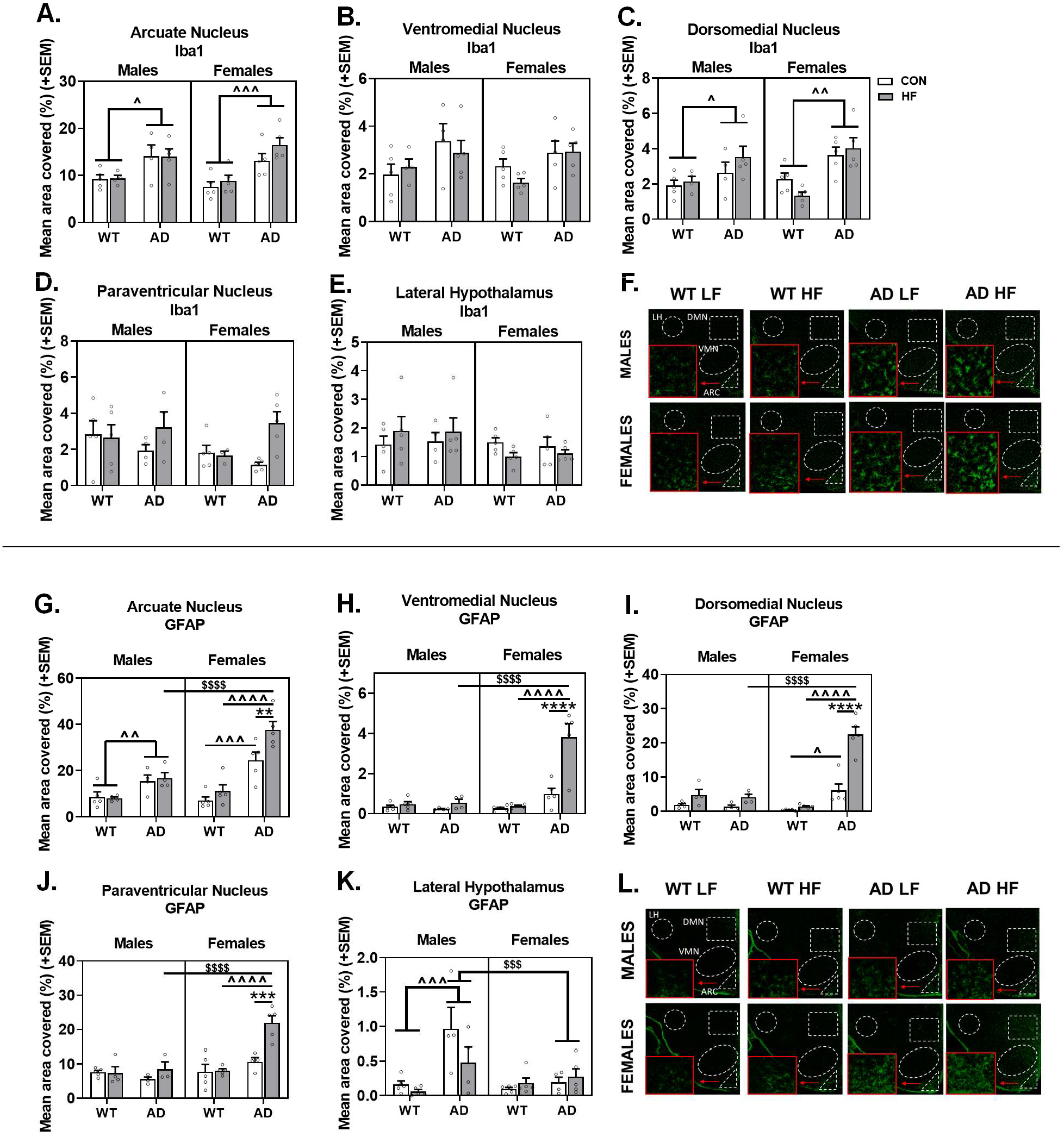
Sex differences in hypothalamic gliosis in response to AD and HF diet. Labeling of tissue sections including the hypothalamus was performed for Iba1 (A-F) and GFAP (G-L). Quantification was performed for % area positive for each label in the arcuate nucleus (A,G), ventromedial nucleus (B,H), dorsomedial nucleus (C,I), paraventricular nucleus (D,J), and lateral hypothalamus area (E,K). Representative images for labeling with Iba1 (F) and GFAP (L) are also shown. * = diet effect p<0.05; ** = diet effect p<0.01; *** = diet effect p<0.001; **** = diet effect p<0.0001 $ = sex effect p<0.05; $$ = sex effect p<0.01; $$$ = sex effect p<0.001; $$$$ = sex effect p<0.0001 ^ = AD effect p<0.05; ^^ = AD effect p<0.01; ^^^ = AD effect p<0.001; ^^^^ = AD effect p<0.0001

Correlations were run to test the hypothesis that hypothalamic microgliosis was associated with metabolic disturbances **(Table 1)**, and correlations were seen in the PVN only and was restricted to AD mice. In both male and female AD mice, Iba1 labeling in the PVN was positively associated with weight gain, though this reached statistical significance in females only.

#### GFAP labeling

Immunolabeling was performed for GFAP to localize astrogliosis in hypothalamic nuclei that regulate energy balance **(Figure 5G-L)**. In the arcuate nucleus, HF diet resulted in an increase in reactive astrocytes in females but not males (p=0.002). AD mice had greater area covered by GFAP-positive label compared to WT mice, in both males (p=0.009) and females (p<0.001). The HF diet-induced increase in GFAP was greater in AD females compared to males (p<0.001), with AD HF females having the greatest GFAP coverage of all groups.

Trends of GFAP labeling were similar within the DMH, VMH, and PVN. In these regions, there was a main effect of HF diet having greater coverage than control diet (p<0.01 for all). Additionally, AD females had greater GFAP labeling than both WT females and AD males (AD x sex interaction p<0.0001). The most striking effect was seen in AD HF females, having ~3-4x or more coverage than all other groups in these three regions (p<0.0001 for all).

There was a clearly different trend in the LHA, with a significant AD x sex interaction (p=0.0089) showing that AD males had greater GFAP coverage compared to WT males and AD females (p≤0.001 for both).

Correlations were run to test the hypothesis that hypothalamic astrogliosis was associated with metabolic disturbances **(Table 1)**. Positive associations were seen between GFAP labeling in the DMH and weight gain in all groups (WT male, AD male, WT female, AD female). However, only in AD females was weight gain positively associated with GFAP labeling in the ARC, VMH, and PVN.

## Discussion

Metabolic abnormalities are associated with increased risk of AD, which may be both a result of and contribute to AD pathology, hypothalamic dysfunction, and behavioral deficits [2]. In the current study, we investigated the interaction between AD and HF diet to influence peripheral and hypothalamic measures of metabolism and inflammation in male and female 3xTg-AD mice. Exploring sex differences in these relationships was of interest given the sex bias of AD (females more greatly affected than males) [49] and to determine whether the potential utility of any findings as novel biomarkers or targets for treatment may be sex-specific. Here, we report that AD males and females exhibit abnormalities in body/fat mass and glucose intolerance in opposite directions (increased in females, decreased in males). AD males exhibit fairly widespread increases in peripheral and hypothalamic inflammation, with the former exacerbated by HF diet in some cases. HF diet in AD females results in severe astrogliosis and increased IL-1B expression in the hypothalamus. These findings may represent two metabolic phenotypes (weight loss and obesity/weight gain) seen in AD patients, with different physiological and/or neurobiological mechanisms.

Here, we observed drastic sex differences in metabolic status of 3xTg-AD mice on a control diet relative to WT mice. Adult male 3xTg-AD mice exhibit decreased body weight and fat mass and better glucose tolerance compared to their WT counterparts. This is in line with clinical findings that report unintentional weight loss in up to ~40% of cases [6]. Conversely, AD females exhibited metabolic disturbances opposite to males, such that AD females displayed increased body and fat mass, in addition to impaired glucose tolerance. This is in line with AD patients having heightened risk for metabolic disease such as T2D [37]. Additionally, in mice ICV injections of Aβ oligomers leads to peripheral glucose intolerance [48]. Little data is available on sex differences in humans, though one study reported that both male and female AD patients exhibited significant weight loss (−2.5kg/year) compared to healthy controls [83]. Our findings are also in contrast to a previous study in Tg2576 mice, which found similar deficits in body mass and fat in males and females [24]. Of note, in both sexes, altered body mass (decreased in males and increased in females) in our AD mice was not observed at the start of the experiment (3-4 months of age), suggesting metabolic disturbances progress with disease status. Although clinically important weight *loss* is more common in AD patients compared to normal older controls, this same study also found that significant weight gain (>5%) was more likely to occur in AD patients [84], suggesting that two metabolic phenotypes may exist in AD patients, which may be represented by our male and female mice.

It has been hypothesized that weight loss may be in part explained by reduced food intake attributable to changes in appetite, impaired autonomy, or cognitive-related issues [85]; however, studies have found that AD patients often exhibit normal or even increased food consumption [86–88]. In fact, concomitant overeating and weight loss has been observed in AD [89]. Another study reported that both male and female AD patients similarly exhibited weight loss versus healthy controls, despite all groups eating a standard institutionalized diet and exhibiting similar energy expenditure [83]. In agreement with this patient data, we saw no differences in caloric intake between 3xTg-AD and WT males. Other AD rodent models similarly report body weight deficits that become more pronounced with age, as well as decreased body fat, despite similar or even increased food consumption [24, 90–94]. While unclear in clinical populations, findings in mice exhibiting AD pathology suggest that elevated basal metabolic rate (measured by oxygen consumption), rather than increased activity levels, may be contributing to these differences [24, 91, 92, 95]. There are some studies that do show hyperactivity in AD mouse models [93]; however, AD males in the current study did not exhibit increased activity levels in open field testing. It should be noted, however, that these were short tests (10 min) and could be reflective of reactivity to a novel environment and influenced by anxiety levels.

In line with low BMI/body mass, hypoleptinemia has been demonstrated in AD patients [96] and mouse models [24], observed prior to AD plaques [24], and with severity becoming more pronounced through the course of the disease. Here, we report nearly non-existent levels of circulating leptin in control-fed AD males, with no changes in hypothalamic expression of LepR, with the latter in agreement with findings in other mouse models [24]. Hypoleptinemia may be due not only to the presence of less adipose tissue, but also attenuated leptin secretion by adipocytes, as seen in mouse models of AD [90]. Impaired leptin signaling is not only believed to be a result of AD pathology, but it may also further promote AD pathology and cognitive deficits. In AD mouse models, low leptin is associated with impaired cognitive function and increased Aβ burden [97–99]. Clinical studies have shown that hypoleptinemia is associated with decreased hippocampal gray matter [100] and cognitive decline in older adults [101]. Leptin has been explicitly shown to play a role in maintaining proper structure and function of the hippocampus [102] and is protective against Aβ and tau pathology [27]. Taken together, these findings support a growing body of evidence that normalizing metabolic dysfunction and perhaps specifically leptin signaling could prove to be a novel target for treatment of AD [31].

It appears that control-fed male AD mice exhibit some appropriate metabolic responses to having reduced adiposity and low circulating leptin, such as increased plasma ghrelin. In healthy older adults, higher circulating ghrelin levels are associated with poorer cognitive performance across several domains [103], while serum levels of the activated (acylated) form of ghrelin were significantly elevated in MCI patients [104]. Therefore, altered ghrelin levels appear to precede cognitive impairment in AD, though there is little to no evidence to support that this may contribute to disease course. In fact, ghrelin is believed to have neuroprotective effects, with ghrelin and agonists alleviating amyloid pathology, boosting neurogenesis, and improving cognition in mouse models of AD [105–107]. In addition to increased ghrelin, control-fed AD males also exhibit increased hypothalamic expression of NPY and AgRP. These changes may represent an inadequate compensatory mechanism to increase caloric intake and reduce energy expenditure to restore body fat and circulating leptin levels. In female Tg2576 mice (amyloid only model), reduced body mass/adiposity was associated with reduced plasma leptin and appropriate reductions in POMC and CART, while NPY and AgRP were unchanged [24]. However, ICV injection of Aβ oligomers into WT mice resulted in hypothalamic amyloid accumulation and increased expression of AgRP and NPY, with no change in POMC [48]. This was suggested to be mediated by Aβ oligomer-induced microglial activation and release of TNF-α to increase production of neuronal AgRP and NPY [48]. In addition to altered plasma levels and hypothalamic expression, it is possible that hypothalamic neurons are less responsive to metabolic signals in AD [24]. Given the overlap of vascular disease and AD, it is also worth noting that vascular dysfunction could also contribute to impaired metabolic function in dementia, since the ability of peripheral signals to reach the hypothalamus is dependent on proper blood flow. In fact, non-significant trends of decreased hypothalamic perfusion have been demonstrated using SPECT in AD patients [19, 20].

Few studies have assessed sex differences in the effects of HF diet in AD mice, despite the sex bias in AD [49], high rates of comorbidity of metabolic disease and AD in humans [37], and that HF diet (particularly saturated fat) and metabolic dysfunction are believed to contribute to AD risk and progression [35, 36]. Here, we report that WT and AD males had similar weight and glucose intolerance on HF diet; however, AD females were considerably more susceptible to increases in body and fat mass, as well as glucose intolerance, in response to HF diet compared to all other groups. These findings provide additional evidence for increased susceptibility to diet-induced metabolic dysregulation in the presence of AD pathology [108]. Of note, AD HF fed females displayed approximately 2-3x as much visceral fat compared to all other groups on the same diet. Visceral fat is thought to be the “bad” fat, contributing to inflammation, disease risk (increased risk of diabetes, CVD, stroke), and linked to cognitive deficits and dementia risk above and beyond obesity [109]. In line with previous studies, HF diet increased TNF-α in adipose tissue likely due to macrophage infiltration [110]; however, this was only observed in AD mice. Additionally, only in males did AD mice exhibit increased IL-1β expression in visceral fat. These findings support a previous study demonstrating that mice expressing mutant APP exhibit altered adipose biology under both obesogenic and non-obesogenic conditions linked to endocrine and metabolic dysfunction [111]. In line with increased adiposity, AD HF females also exhibited hyperleptinemia compared to other female groups, which may have contributed to glucose intolerance. AD HF females also exhibited increased heart mass, likely to support blood flow throughout a body with increased mass; this is in line with previous findings suggesting that HF diet/obesity are associated with increased cardiac stress [112, 113]. While differences in energy intake or output (activity levels) could not explain findings in AD females on control diet, AD females on HF diet consumed a greater number of calories and were hypoactive, which contribute to an energy surplus state. Our 3xTg-AD model may be a model of “accelerated aging”, in which case our findings are in line with our previous study reporting that female mice exhibit increased weight gain and glucose intolerance compared to males in middle age on HF diet (~15mo) [53].

There is ample evidence to suggest that there is an interrelationship between nonalcoholic fatty liver disease (NAFLD) and metabolic disease. High fat diet and increased adiposity can cause an increase in fatty acids leading to hepatic steatosis. Fat accumulation in the liver can lead to lipotoxicity and inflammation, causing oxidative damage, ER stress, and mitochondrial dysfunction, which in turn causes hepatic insulin resistance, de novo lipogenesis, and later peripheral insulin resistance due to hyperglycemia and dyslipidemia [59]. It is possible that increased macrovesicular fat in AD females on control diet could have contributed to their glucose intolerance, as liver fat content shows greater association with glucose intolerance than even visceral fat [114]. In the current study, AD males and females accumulated similar amounts of microvesicular and macrovesicular fat in the liver in response to high fat diet, though AD mice overall had greater macrovesicular fat and hepatic fibrosis. Of note, AD males had greater leukocyte accumulation, a sign of potential inflammation in the liver, compared to other groups; this was exacerbated by HF diet. Increased leukocyte accumulation in the liver in response to HF diet was also seen in AD females, albeit to a lesser degree. These findings were somewhat surprising, given results on weight gain, adiposity, and glucose tolerance in these groups. However, more severe NAFLD pathology in AD males on HF diet was in line with findings that several other plasma markers associated with diabetes were disproportionately increased in this group, despite adiposity that was less than or equal to other HF-fed groups. Despite AD HF females exhibiting increased weight gain, adiposity, and glucose intolerance compared to all other groups, many other metabolic markers in circulation or expression in the hypothalamus were not seemingly disproportionately altered. This is in line with a previous study in 3xTg-AD mice showing males but not females exhibit hyperinsulinemia in response to HF diet; however, these males also displayed increased ectopic fat despite similar weight gain and hyperglycemia, in contrast to current findings [43].

NAFLD and other chronic liver diseases are associated with increased dementia risk, even in the absence of metabolic disease such as Type 2 diabetes [115]. NAFLD is associated with altered cerebral blood flow, increased risk and severity of stroke, asymptomatic brain lesions, brain aging, and cognitive impairment [61]. Additionally, the healthy liver plays a role in the peripheral metabolism of amyloid; therefore, in cases of liver disease like NAFLD, impaired peripheral metabolism can reduce the efflux of AB from the brain [61]. In fact, animal models of NAFLD show enhanced amyloid and tau pathology, as well as neuroinflammation [116, 117]. Additionally, NAFLD is characterized by a pro-inflammatory state, which may at least in part be linked to the increased systemic inflammation (plasma IL-10, IL-12, MIP-1a, and MIP-1b) seen in AD males and exacerbated by HF diet in this group, mimicking trends of increased leukocyte accumulation in the liver.

Both AD and metabolic disease are linked to chronic systemic inflammation, which can lead to tissue damage and impaired function. Specifically, hypothalamic inflammation in obesity is thought to contribute to impaired action of leptin, insulin, and other hormones [118]. In the current study, increased inflammation in the periphery of male AD mice (liver and plasma) mimics trends of increased hypothalamic expression of Iba1 and GFAP, and pro-inflammatory cytokines (TNF-a and IL-1B). In the current study, increased inflammation in the periphery of AD mice (liver, plasma, and visceral fat) mimics trends of increased hypothalamic expression of glial markers Iba1 and GFAP, as well as pro-inflammatory cytokines (TNF-a and IL-1B), in AD males. Previous studies have also reported increased hypothalamic expression of pro-inflammatory markers in 3xTg-AD male mice at even 12 weeks of age [95]. Additionally, free fatty acids from the consumption of a high fat diet can contribute to hypothalamic inflammation [119]. Inflammation and impaired liver function are hypothesized to contribute to BBB breakdown, which would allow for the infiltration of peripheral immune cells [61]. Of note, the purposeful leakiness of the blood brain barrier in the mediobasal hypothalamus, which normally functions to allow the sensing of peripheral signals, also leaves it particularly vulnerable to this source of inflammation. Additionally, microglia are targets of circulating cytokines, including liver-derived metabolic inflammation, promoting further neuroinflammation through the release of proinflammatory cytokines [120]. NAFLD induced by administration of CCl_4_ results in significant increases in brain levels of TNF-a and IL-6, as well as neurodegeneration [121]. Immunohistochemical analyses also revealed that AD males exhibited microgliosis and astrogliosis in several hypothalamic nuclei involved in energy balance, a phenomenon that was largely unaffected further by HF diet. In contrast to AD males, AD females do not exhibit significant increases in plasma markers of inflammation, even on HF diet. AD HF females did exhibit moderately elevated liver inflammation, as well as hypothalamic expression of only IL-1B; however, this was to an equal or lesser degree compared to male counterparts. Our findings may be reflective of theories that high-grade inflammation is associated with unintentional weight loss (as in control AD males, counteracted by HF diet) while low-grade inflammation is associated with obesity/metabolic disease [118, 122].

In addition to contributing to inflammation under obesigenic conditions, previous studies have demonstrated that glial cells play a multifaceted role in energy balance, potentially in a sex-specific manner [123]. The most striking difference in the hypothalamus of AD females, particularly on a HF diet, was a marked increase in GFAP expression in the whole hypothalamus, as well as increased GFAP labeling in the arcuate nucleus (ARC), dorsomedial hypothalamus (DMH), ventromedial hypothalamus (VMH), and paraventricular nucleus (PVN). Although GFAP staining in the DMH was correlated with weight gain in all groups, staining in the ARC, VMH, and PVN, as well as whole hypothalamus expression of GFAP, were positively correlated with metabolic outcomes such as weight gain in AD females only. These findings suggest that the metabolic deficits in AD females and enhanced susceptibility to HF diet may be at least in part attributed to astrogliosis in these hypothalamic nuclei. Astrocytes can become reactive in response to leptin and IL-1β, both of which were increased in AD females. This proposed mechanism is supported by findings that both plasma leptin and hypothalamic IL-1β expression were positively correlated with hypothalamic expression of GFAP, and IL-1β expression was associated with metabolic outcomes, in AD females only. Additionally, In addition to releasing pro-inflammatory mediators, activation of astrocytes by cytokines such as IL-1β can result in NMDA receptor-dependent neuronal death, increased extracellular glutamate, and hyperexcitable synaptic activity [124]. Additionally, astrocytes orchestrate synaptic remodeling and are capable of ensheathing neurons in the arcuate nucleus, rendering them less responsive to metabolic signaling [125]. In AD HF females, this could contribute to their inability to sense signals trying to maintain energy and glucose homeostasis. In line with increased adiposity, AD HF females exhibited hyperleptinemia; however, they did not demonstrate a downregulation of LepR in the hypothalamus as might be expected; instead, there was a trend of increased LepR expression that correlated with weight gain in AD females. Although previous studies suggest that amyloid is capable of attenuating hypothalamic neuron reactivity to leptin and ghrelin [24], evidence from both mouse models and AD patients suggest that hypothalamic impairment and metabolic abnormalities precede amyloid and tau pathology [15, 126]; therefore, other mechanisms must be playing a role. Astrogliosis and the release of pro-inflammatory cytokines may be contributing to the two metabolic phenotypes reflected in male and female 3xTg-AD mice in the current study; however, further investigation is necessary to confirm and elucidate these possible mechanisms.

Extra-hypothalamic brain regions that could also be contributing to sex- and diet-dependent metabolic differences in AD mice have been comparatively underexplored. The nucleus of the solitary tract in the brain stem senses peripheral metabolic signals and sends and receives projections to/from the hypothalamus to regulate energy balance [81, 127, 128]. Function of the nucleus of the solitary tract has been shown to be impaired by both high fat diet and is hypothesized to play a role in AD [129, 130]. In addition to the homeostatic regulation of feeding behavior, the brain’s reward system, including the mesolimbic and mesocortical dopaminergic pathways, controls hedonic feeding to promote the consumption of palatable foods and can override satiety signals. Dopaminergic neurons in the ventral tegmental area appear to be compromised in both AD mice and late-onset patients [131–133]. In AD mice, this is associated with attenuated dopaminergic output in the hippocampus and nucleus accumbens, as well as altered neural and behavioral reactivity to reward, including palatable foods [131, 134]. Additionally, high fat diet and obesity can also contribute to altered dopaminergic signaling in the reward pathway [135], possibly-compounding AD-related dysfunction. Further exploration of the possible contribution of these regions to altered metabolic outcomes could yield additional targets for treatment.

## Conclusions

In summary, we found marked sex differences in metabolic outcomes of AD mice both on control and a HF diet, which may be representative of two metabolic phenotypes associated with Alzheimer’s disease. We found that on control diet, AD males and females exhibit metabolic changes in opposite directions, with males representing an energy deficit state unattributable to energy intake/output, and females an energy surplus. Our results confirm findings that both weight loss/low body mass index and weight gain/glucose intolerance may be possible biomarkers for AD. Hypothalamic inflammation in control fed AD mice was worse in males than females, in line with some increases in markers of peripheral inflammation. In males, we see a somewhat “normal” metabolic response to HF diet and no additive effect on neuroinflammation, though there is a clear exacerbation of systemic inflammation and some peripheral markers of metabolic disease. In AD females, we see a dramatic increase in susceptibility to some metabolic effects of HF diet (weight gain, adiposity, glucose intolerance) associated with increased caloric intake and decreased energy expenditure, as well as hyperleptinemia, that is associated with severe astrogliosis in the hypothalamus. Further exploration is necessary to elucidate these potential mechanisms and determine possible new targets for treatment.

## List of abbreviations

α-MSH: Alpha-melanocyte-stimulating hormone
Aβ: Amyloid-beta
AD: Alzheimer’s disease
AgRP: Agouti-related peptide
ANOVA: Analysis of variance
ARC: Arcuate nucleus
AUC: Area under the curve
BMI: Body mass index
CART: cocaine- and amphetamine-regulated transcript CON; Control (diet)
CSF: Cerebrospinal fluid
DMH: Dorsomedial hypothalamus
EDTA: Ethylenediaminetetraacetic acid
FNDC5: Fibronectin type III domain-containing protein 5
GFAP: Glial fibrillary acidic protein
GIP: Gastric inhibitory polypeptide
GLP-1: Glucagon-like peptide-1
GTT: Glucose tolerance test
H & E: Hematoxylin and eosin
HF: High fat (diet)
Iba1: Ionized calcium binding adaptor molecule 1
IL-1β: Interleukin-1 beta
IL-6: Interleukin-6
IL-10: Interleukin-10
IL-12 (p40): Interleukin-12 (p40)
LepR: Leptin receptor
LHA: Lateral hypothalamus area
MCR4: melanocortin receptor 4
MIP: Macrophage inflammatory protein
NAFLD: Nonalcoholic fatty liver disease
NPY: Neuropeptide Y
PBS: Phosphate saline buffer
POMC: Pro-opiomelanocortin
PVN: Paraventricular nucleus
SEM: Standard error of the mean
T2D: Type 2 diabetes
TNF-α: Tumor necrosis factor alpha
VMH: Ventromedial hypothalamus
WT: Wild-type

## Declarations

### Ethics approval and consent to participate

Not applicable

### Consent for publication

Not applicable

### Availability of data and materials

The datasets used and/or analyzed during the current study are available from the corresponding author on reasonable request.

### Authors’ contributions

KLZ obtained funding for the experiments. LSR, MAT, and KLZ designed the experiments. OJG, MAT, and AES performed the animal work. LSR, OJG, MAT, AES, and CAG performed the experiments. LSR, OJG, and MAT analyzed the data. LSR and MAT prepared the figures. LSR prepared the manuscript. OJG and KLZ edited the manuscript. All authors approved the final manuscript.

## Acknowledgements

The authors would like to thank Dr. Alexander Mongin for their technical guidance on qPCR experiments. Thank you to Febronia Mansour, Nathan Albert, Alvira Tyagi, and Allegra Wu for assistance with tissue collection and imaging, and Molly Hyer for advice on analysis of liver pathology.

